# Evaluation of azithromycin or hydroxychloroquine plus azithromycin combination therapy on cardiac conduction and function in guinea pigs

**DOI:** 10.1101/2020.10.31.362566

**Authors:** Xiang Li, Weijiang Tan, Shuang Zheng, Huan Sun, Xiaoshen Zhang, Xiaohui Li, Honghua Chen, Xuecong Ren, Tianzhen He, Caiyi Zhu, Yu Zhang, Feng Hua Yang

## Abstract

**Background:** In the early stages of the coronavirus disease pandemic, the anti-malarial drug hydroxychloroquine (HCQ) and the antibiotic drug azithromycin (AZM) were widely used as emerging treatments. However, controversial cardiac toxicity results obtained from clinical trials and epidemic studies suggest that the cardiotoxicity of these two drugs should be re-evaluated. In the present study, we aimed to assess the impact of a short course of AZM or HCQ + AZM combination treatment on ECG and cardiac function in healthy guinea pigs.

**Methods:** Thirty-two male guinea pigs were randomly divided into four groups: control; AZM; HCQ; and HCQ + AZM groups. At 3, 6, and 9 days after treatment, electrocardiograms (ECGs) and echocardiographic techniques were used to determine important ECG parameters and cardiac functional parameters of the left ventricle (including posterior wall thickness, end systolic/end diastolic volume, ejection fraction, and fractional shortening).

**Results:** Although AZM decreased the heart rates of guinea pigs on day 9 (under anesthetized conditions), HCQ + AZM decreased heart rates on days 3, 6, and 9. The corrected QT intervals of guinea pigs after AZM and HCQ + AZM treatments were significantly increased, compared with CON and HCQ treatment respectively, on days 3, 6, and 9. However, QRS complex durations were not significantly different between the groups. AZM significantly decreased left ventricular ejection fraction (LVEF) and left ventricular fraction shortening (LVFS) on days 3, 6, and 9, whereas HCQ + AZM only decreased LVEF and LVFS on day 9. Posterior wall thickness and of the left ventricle in the diastolic and systolic states were not significantly different between these groups. In addition, compared with CON, AZM and HCQ decreased the EDV. And, in comparison with HCQ treatment, HCQ + AZM treatment increased ESV on day 9.

**Conclusions:** According to our study, AZM significantly prolongs the QT interval and damages cardiac function. Moreover, HCQ + AZM treatment increased the risk of cardiac dysfunction compared with HCQ treatment.

## Introduction

Since the onset of the coronavirus disease (COVID-19) pandemic, confirmed cases of COVID-19 have reached more than 44.5 million, and ~2.64% of these cases have proved fatal (WHO Coronavirus Disease (COVID-19) Dashboard, accessed Oct 30, 2020)(26). Hydroxychloroquine (HCQ), azithromycin (AZM), or a combination of these two drugs has been widely used for the treatment of COVID-19, despite that some countries have declined treatment with these drugs. HCQ is a quinoline medicine that was first approved by the FDA in 1955 and has since been widely used for the treatment of malaria, rheumatoid arthritis, and systemic lupous erythematosus(3, 12). AZM is a macrolide antibiotic that was discovered by the Croatia-based Pliva Pharmaceutical Co. in 1988. AZM is used to treat bacterial infections of the respiratory tract, urogenital system, connective tissues, and other systemic infections. Although the early data suggested that treatment with HCQ or AZM reduced the viral load and/ or improved clinical conditions(24), several cohort studies and clinical trials concluded that HCQ monotherapy or HCQ + AZM combination therapy did not improve clinical status (4, 16). Besides of this argument regarding the treatment clinical outcomes, these controversial data concern the safety profiles of these drugs.

According to statistics provided by @CovidAnalysis(1), patients in many countries still use HCQ to treat COVID-19 in the early stages of treatment. HCQ and AZM are currently in the World Health Organization’s list of essential medicines and, even before the pandemic, were among the most commonly prescribed drugs worldwide(6, 7, 27). So far, clinicians and researchers globally have been desperately sharing their findings and experiences regarding the prevention and treatment of COVID-19. However, rapid publication of these findings has also exposed the public to incompletely analyzed, un-verified data. In the assessment of drug-induced cardiotoxicity risk for novel pharmaceuticals, since 2005 most countries have adopted the standard preclinical evaluation protocol of the International Conference on Harmonisation of Technical Requirements for Registration of Pharmaceuticals for Human Use (ICH S7B) (13). However, the drugs including HCQ and AZM on market earlier than that timepoint might not be assessed adequately for their hidden cardiotoxicity. To further understand the adverse effects of HCQ, AZM, and HCQ + AZM on cardiac conduction and function and to validate safe use of these drugs in non-COVID-19 patients, a re-evaluation of cardiac safety using a preclinical animal model is necessary.

Here, we firstly tested delayed ventricular repolarization in guinea pigs after the drug treatment following the guideline of ICH S7B. Next, we also employed echocardiography technology for small animals to evaluate cardiac morphology and function that were potentially affected by drugs.

## Materials and methods

### Animals

Healthy Hartley guinea pigs (~300 g) were purchased from a licensed laboratory animal supplier (Guangdong Medical Laboratory Animal Center, China). The animals were housed in a specific pathogen-free, AAALAC-accredited facility of Guangdong Laboratory Animals Monitoring Institute (Guangzhou, China). The facility employed a 12-h light/dark cycle and a temperature and humidity of 24 ± 2°C and 40-60%, respectively. Animals were fed ad libitum on a standard guinea pig diet. All animal experimental protocols were approved by the Institutional Animal Care and Use Committee of the Guangdong Laboratory Animals Monitoring Institute, Guangzhou, China (No. IACUC2020125).

### Treatment protocol

Thirty-two guinea pigs were randomly divided into four groups (n = 8 in each group): control (CON); azithromycin (AZM); hydroxychloroquine sulfate (HCQ); and HCQ plus AZM (HCQ + AZM) groups. The dosage of drugs was adjusted with reference to the clinical dosage in humans(8), and all drugs were suspended in normal saline before administration. HCQ was purchased from Shanghai Pharmaceuticals (Shanghai, China) and AZM from Pfizer Pharmaceuticals (New York, USA). The experimental design is shown in Figure 1. Specifically, the drugs, dosage, and duration of administration of each group were as follows: (1) CON group, normal saline; (2) HCQ group, 30.84 mg/kg of HCQ on the first day and 15.42 mg/kg of HCQ on subsequent 8 days; (3) AZM group, 38.54 mg/kg of AZM on the first day and 19.27 mg/kg from day 2-5; (4) HCQ + AZM group, 30.84 mg/kg of HCQ on day 1 followed by 15.42 mg/kg per day for the next 8 days and 38.54 mg/kg of AZM on the first day and 19.27 mg/kg from days 2-5. All drugs or normal saline were administered once a day by gavage. Electrocardiogram (ECG) and echocardiograph recordings of guinea pigs were performed before initiation of the administration protocol and on days 3, 6, and 9 after the administration of drugs.

**Figure 1.**
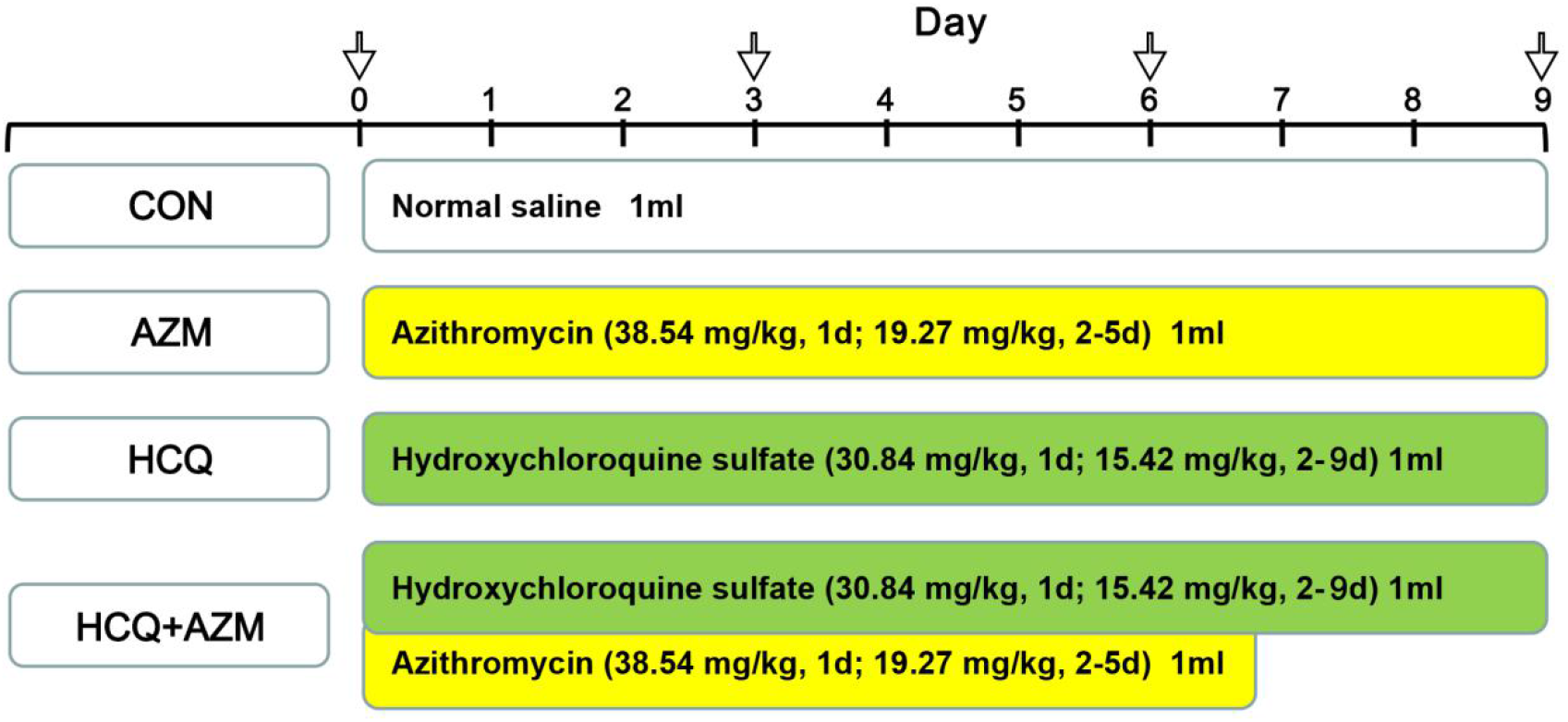
Sketch of dosing regimens and detection point. Arrows indicate time points for electrocardiogram and echocardiographic detection. CON, control group; AZM, Azithromycin group; HCQ, Hydroxychloroquine sulfate group; HCQ + AZM, Hydroxychloroquine sulfate + Azithromycin group.

### *In vivo* electrophysiological parameters

Delayed ventricular repolarization is reflected in QT interval prolongation, which can cause a potentially lethal arrhythmia called Torsade de pointes(14). Longitudinal studies have shown that QT prolongation can help predict cardiovascular event-related mortality(22, 23). To determine the QT interval and other in vivo electrophysiological parameters, the guinea pigs were anesthetized with 2% isoflurane and pure oxygen mix through a respiratory system (Matrx, USA). After induction of anesthesia, the animals were placed in dorsal recumbency on an operating table with a heating pad. Sterilized electrodes were inserted subcutaneously into the right forelimb and the left hind limb of the anesthetized animal. The electrodes were then connected to the ECG module of the PowerLab 4/35 system (ADInstruments Inc., USA), and the lead II method was selected from the program for ECG recording. Electrophysiological parameters, including RR interval, PR interval, QRS duration, and QT interval, were analyzed using the LabChart Pro Software (ADInstruments Inc., USA).

### QT interval correction

Several formulas have been developed for QT interval correction to facilitate a precise interpretation of this interval. In our current experiments, two steps were performed to determine the correction formula most appropriate for calculating the QT interval for the anesthetized guinea pigs. First, the QT and RR intervals from the ECGs were inputted into five correction equations (Bazett’s(2), Fridericia’s(11), Van de Water’s (25), Kawataki’s(15), Matsunaga’s(17) to obtain the corrected QT (QTc) interval. Next, the calculated QTc and RR data were subjected to linear regression analysis, and the correction formula with the smallest slope of the regression line was used to evaluate the QTc interval in this study.

1. Bazett: QTc = QT/RR1/2
2. Fridericia: QTc = QT/RR1/3
3. Van de Water’s: QTc = QT – 0.087 (RR – 1000) = QT – 87 (60/HR – 1)
4. Kawataki: QTc = QT/RR1/4
5. Matsunaga: QTc = log600 × QT/logRR

### Transthoracic echocardiography

The effects of drugs on the cardiac function of guinea pigs were evaluated using a high-frequency ultrasound system Vevo2100 (VisualSonics, Canada) equipped with a linear array transducer (MS250, 13-24 MHz). This transducer is specifically designed for rats, guinea pigs or other similar sizes of small animals. During image acquisition, guinea pigs were anesthetized continuously with 2% isoflurane. All animals were placed in the dorsal recumbent position for ultrasound imaging. The probe was first placed on the right shoulder approximately 30° from the center line to obtain the parasternal left ventricle long axis B-mode image, and the probe was then rotated 90° clockwise to obtain the parasternal short-axis B-mode image of the left ventricle on the papillary muscle plane. The M-mode ultrasound cursor was used to acquire images.

### Cardiac function parameters

A VisualSonics workstation was used to analyze the echocardiographic images. Using the short-axis M-mode images, the left ventricular dimensions (LVIDd and LVIDs), and posterior wall thickness (LVPWd and LVPWs) were all measured at the end of both systole and diastole. The end diastolic volume (EDV), end systolic volume (ESV), ejection fraction (LVEF), and fractional shortening (LVFS) of the LV were then calculated. The calculation equations for EF and FS were:

1. EF = (EDV – ESV)/EDV
2. FS = (LVIDd–LVIDs)/LVIDd

All ultrasound data were averaged over three consecutive cardiac cycles.

### Statistical Analysis

All experimental data are presented as means ± standard error of means (SEM). Statistical differences between the groups were analyzed by two-way ANOVA followed by Tukey’s multiple-comparisons test (GraphPad Prism 8, USA). P < 0.05 was considered statistical significance.

## Results

### Evaluation of QT correction methods for estimating QT interval prolongation

The correlation (R^2^) values obtained using the QT interval correction formulas reported by Bazett, Van de Water, Kawataki, Matsunaga, and Fridericia were 0.3446, 0.6661, 0.5871, 0.6360, and 0.5179, respectively (Figure 2). Thus, the Van de Water, Kawataki, and Matsunaga methods yielded the highest correlations. In a comparison of the slopes obtained from data fitted to these three linear regression equations, the smallest value (0.4947) was returned using the Van de Water equation (Figure 2, Table 1). Thus, the Van de Water correction method was used to evaluate the QT interval in this study.

**Figure 2.**
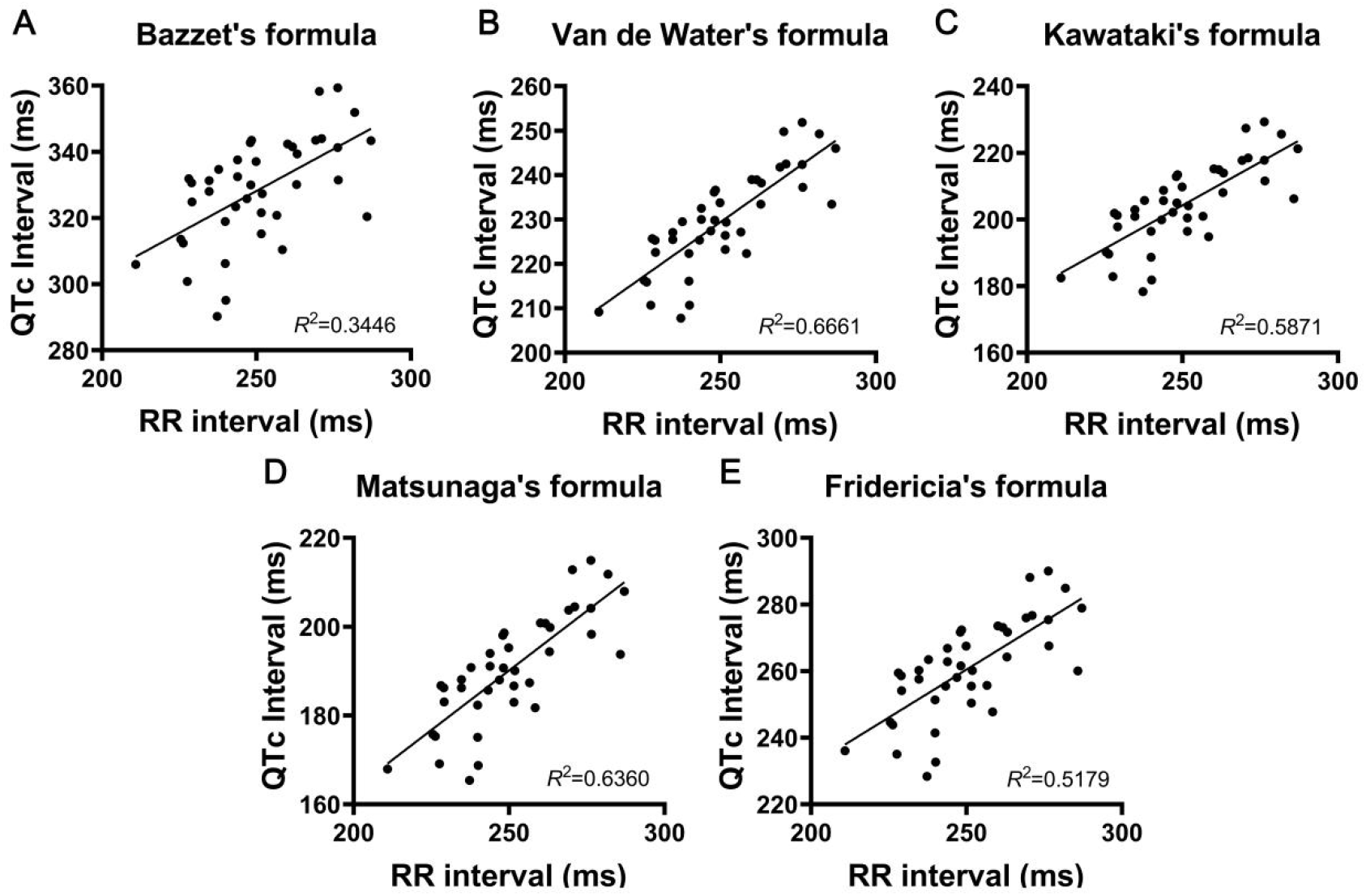
Scatter plots used for linear regression between corrected QT interval (QTc) and RR interval in guinea pigs under anesthesia. Data were collected from 40 health guinea pigs. Correction formulas including Bazett’s (A), Van de Water’s (B), Kawataki’s (C), Matsunaga’s (D), and Fridericia’s (E).

**Table 1.** QT correction formulas assessed in this work. Data were collected from 40 health guinea pigs. RR, RR interval; QTc, Corrected QT.

| Name | Formula | Linear regression |  |  |
| --- | --- | --- | --- | --- |
|  |  | Slope | Intercept | R<sup>2</sup> |
| Bazzet | $QTc = QT \times (1000/RR)^{1/2}$ | 0.5073 | 201.3 | 0.3446 |
| Van de Water | $QTc = QT - 0.087(RR - 1000)$ | 0.4947 | 105.7 | 0.6661 |
| Kawataki | $QTc = QT \times (600/RR)^{1/4}$ | 0.5196 | 74.34 | 0.5871 |
| Matsunaga | $QTc = QT \times \log 600 / \log RR$ | 0.5361 | 56.09 | 0.6360 |
| Fridericia | $QTc = QT \times (1000/RR)^{1/3}$ | 0.5759 | 116.5 | 0.5179 |

### QTc intervals of Guinea pigs significantly increased with AZM or HCQ + AZM treatment

Sample ECGs on Day 0 and Day 9 from the CON, AZM, HCQ, and HCQ + AZM groups are shown in Figure 3. Basic electrocardiogram (ECG) parameters of guinea pigs before drug administration were shown in Table 2. Although guinea pig heart rates in the HCQ group were unchanged (compared with those in the CON group) on days 3, 6, and 9. Compared with HCQ treatment alone, HCQ + AZM treatment significantly reduced the heart rate on days 3, 6, and 9 (Figure 4A and 4B). Thus, guinea pig heart rate slowed following the addition of AZM to HCQ treatment. And, bradycardia was still observed in guinea pigs after the discontinuation of AZM treatment for 4 days.

**Figure 3.**
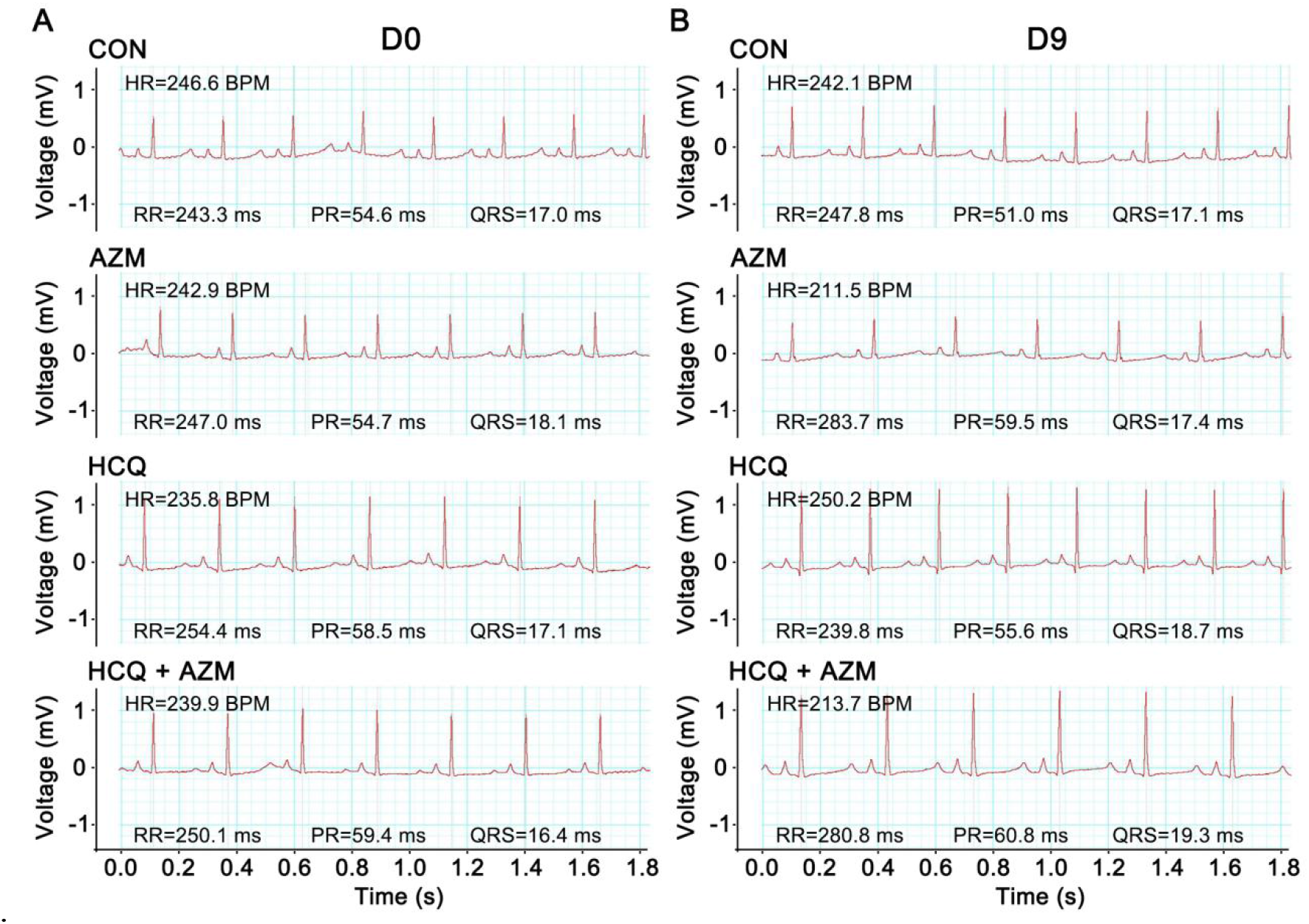
Typical tracings of the surface lead II electrocardiogram *in vivo*. Tracings were obtained before (A) or after 9 days (B) using hydroxychloroquine (HCQ) treatment with or without azithromycin (AZM). ECG, electrocardiogram; CON, control group; AZM, Azithromycin group; HCQ, Hydroxychloroquine sulfate group; HCQ + AZM, Hydroxychloroquine sulfate + Azithromycin group; D, day; HR, heart rate; BPM, beats per minute; RR, RR interval; PR, PR interval; QRS, QRS duration.

**Figure 4.**
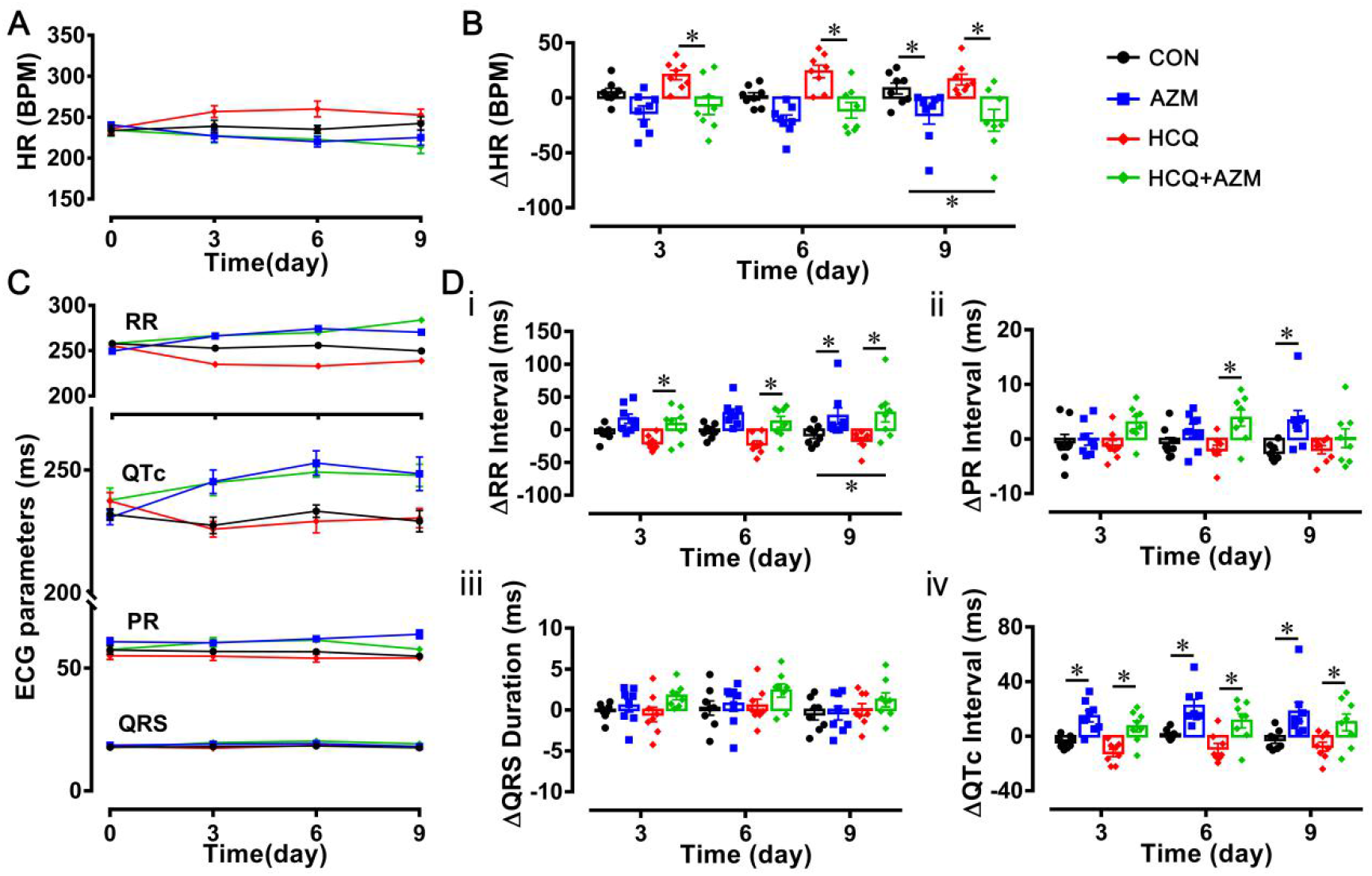
Heart rate (HR) and electrocardiogram (ECG) parameters obtained before drug administration and after 3, 6, and 9 days using hydroxychloroquine (HCQ) treatment with or without azithromycin (AZM). Time courses of changes in HR (A) and ECG parameters (C) before or after the gavage of HCQ with or without AZM. Changes in HR (B) and ECG parameters (D) at each monitoring point after administration were also statistically analyzed. ECG parameters including (i) RR interval; (ii) PR interval; (iii) QRS duration; and (iv) QTc interval. Data are presented as mean ± standard error of mean (n = 8, both groups), *p < 0.05. Statistical significance was determined using two-way analysis of variance coupled with Tukey’s multiple comparison test. ECG, electrocardiogram; CON, control group; AZM, Azithromycin group; HCQ, Hydroxychloroquine sulfate group; HCQ + AZM, Hydroxychloroquine sulfate + Azithromycin group; HR, heart rate; BPM, beats per minute; QTc, Corrected QT.

**Table 2.** Basic electrocardiogram (ECG) parameters of guinea pigs before drug administration. Data are presented as mean ± standard error of mean (n = 8, both groups). Statistical significance was determined using one-way analysis of variance coupled with Tukey’s multiple comparison test. ^a^P< 0.05 vs. CON group; ^b^P<0.05 vs. AZM group; ^c^P < 0.05 vs. HCQ group. ECG, electrocardiogram; CON, control group; AZM, Azithromycin group; HCQ, Hydroxychloroquine sulfate group; HCQ + AZM, Hydroxychloroquine sulfate + Azithromycin group; HR, heart rate; BPM, beats per minute; RR, RR interval; PR, PR interval; QRS, QRS duration; QTc, Corrected QT.

| Group | HR (BPM) | RR (ms) | PR (ms) | QRS (ms) | QTc (ms) |
| --- | --- | --- | --- | --- | --- |
| CON | 233.83 ± 6.33 | 257.93 ± 6.95 | 57.39 ± 1.72 | 18.10 ± 0.60 | 231.91 ± 2.29 |
| AZM | 240.55 ± 3.68 | 249.51 ± 4.08 | 60.41 ± 1.53 | 18.50 ± 0.66 | 230.66 ± 2.97 |
| HCQ | 236.06 ± 6.33 | 255.48 ± 6.96 | 56.05 ± 1.26 | 18.00 ± 0.74 | 238.09 ± 3.29 |
| HCQ + AZM | 234.11 ± 7.49 | 258.09 ± 8.10 | 57.56 ± 1.25 | 17.91 ± 0.28 | 237.73 ± 4.97 |

Correspondingly, the RR interval was significantly increased only on day 9 in the AZM group compared with that in the CON group. The HCQ + AZM treatment significantly increased the RR interval on days 3, 6, and 9 compared with HCQ treatment, while in comparison with CON, HCQ + AZM treatment was significantly increased RR interval on day 9 only (Figure 4C and 4D-i; Table 3).

**Table 3.** Variations in electrocardiogram (ECG) parameters at D0–D3, D0–D6, D0–D9. Δ HR, Δ RR, Δ PR, Δ QRS, and Δ QTc represents differences in respective parameters before and after administration. Data are presented as mean ± standard error of mean (n = 8, both groups). Statistical significance was determined using two-way analysis of variance coupled with Tukey’s multiple comparison test. ^a^P< 0.05 vs. CON group; ^b^P<0.05 vs. AZM group; ^c^P < 0.05 vs. HCQ group. ECG, electrocardiogram; CON, control group; AZM, Azithromycin group; HCQ, Hydroxychloroquine sulfate group; HCQ + AZM, Hydroxychloroquine sulfate + Azithromycin group; D, day; HR, heart rate; BPM, beats per minute; RR, RR interval; PR, PR interval; QRS, QRS duration; QTc, Corrected QT.

| Group | ΔHR (BPM) | | | ΔRR (ms) | | |
| --- | --- | --- | --- | --- | --- | --- |
|  | D0-D3 | D0-D6 | D0-D9 | D0-D3 | D0-D6 | D0-D9 |
| CON | 4.99±3.62 | 1.34±3.32 | 8.50±5.00 | -5.06±3.80 | -2.05±3.77 | -8.15±5.32 |
| AZM | -13.53±6.15 | -20.46±4.82 | -15.53±8.29<sup>a</sup> | 16.75±7.16 | 24.89±6.75 | 20.98±12.38<sup>a</sup> |
| HCQ | 20.71±4.36 | 23.98±5.74 | 16.63±4.86 | -20.48±3.86 | -22.45±5.20 | -16.61±4.96 |
| HCQ + AZM | -6.93±8.29<sup>c</sup> | -11.34±7.14<sup>c</sup> | -20.49±9.93<sup>ac</sup> | 8.59±8.94<sup>c</sup> | 12.10±8.29<sup>c</sup> | 25.78±13.91<sup>ac</sup> |

| ΔPR (ms) | | | ΔQRS (ms) | | | ΔQTc (ms) | | |
| --- | --- | --- | --- | --- | --- | --- | --- | --- |
| D0-D3 | D0-D6 | D0-D9 | D0-D3 | D0-D6 | D0-D9 | D0-D3 | D0-D6 | D0-D9 |
| -0.57±1.41 | -0.72±0.96 | -2.52±0.54 | 0.25±0.45 | 0.61±0.87 | -0.16±0.62 | -4.43±1.58 | 1.36±1.30 | -2.75±2.52 |
| -0.41±1.16 | 1.21±1.32 | 3.02±1.92<sup>a</sup> | 0.43±0.85 | 0.66±0.99 | -0.54±0.91 | 14.61±4.09<sup>a</sup> | 22.19±4.87<sup>a</sup> | 17.83±6.98<sup>a</sup> |
| -0.18±0.96 | -1.00±1.10 | -0.93±0.95 | -1.34±0.99 | 0.53±0.80 | 0.05±0.72 | -12.44±2.85 | -9.00±3.61 | -7.67±3.34 |
| 2.98±1.13 | 3.83±1.49<sup>c</sup> | 0.13±1.73 | 1.98±0.52<sup>c</sup> | 2.63±0.93 | 1.50±0.99 | 7.10±4.05<sup>c</sup> | 11.23±5.14<sup>c</sup> | 10.12±6.13<sup>c</sup> |

The PR interval was significantly increased following a 9-day course of AZM, compared with CON. The PR interval was prolonged on day 6 but not on day 9 following AZM + HCQ treatment, compared with HCQ. These results suggest that conduction of the sinoatrial nodal impulse to the ventricles is affected by AZM and that subsequent withdrawal of AZM halts the observed conduction dysfunction induced by HCQ + AZM treatment on day 6 (Figure 4C and 4D-ii; Table 3). The QRS complex durations were not different among the four treatment groups (Figure 4D-iii; Table 3), indicating that short-course HCQ, AZM, and HCQ + AZM treatments did not significantly affect right or left ventricle depolarization.

Following AZM treatment, the QTc interval was significantly increased on days 3, 6, and 9 compared with that following CON treatment. In contrast, HCQ treatment did not affect the QTc interval on days 3, 6, or 9. However, the QTc interval was significantly increased following HCQ + AZM treatment on days 3, 6, and 9 compared with that following HCQ treatment (Figure 4C and 4D-iv; Table 3). These results suggest that AZM prolongs the QT interval and that after 4 days of AZM withdrawal, QT interval prolongation did not disappear.

### AZM significant decreased LV ejection fraction (LVEF) and LV fractional shortening (LVFS), while HCQ + AZM only decreased LVEF and LVFS on day 9

First, we examined the cardiac morphology of guinea pigs in the CON, AZM, HCQ, and HCQ + AZM groups. From examinations of wall thicknesses and internal dimensions in the diastolic and systolic states, no differences in the LVPWd and LVPWs were observed among the groups on days 3, 6, and 9. However, on day 9, AZM significantly reduced LVIDd compared with CON. Compared with HCQ, HCQ + AZM significantly increased the LVIDs on day 9. And, compared with AZM, HCQ + AZM significantly decreased LVIDs and LVIDd on day 6 but not day 9. These results indicate that short-course treatments of AZM or HCQ + AZM altered the morphology of the LVs. Simultaneously, on day 9, in comparison with CON, the AZM or HCQ treatment alone decreased the EDV, while in comparison with HCQ treatment, HCQ + AZM treatment increased ESV.

Next, we compared cardiac function among the groups (Figure 5; Table 4). In the AZM group, LVEFs were significantly decreased on days 3, 6, and 9 compared with those in the Con group. Furthermore, compared with HCQ treatment, HCQ + AZM treatment for 5 days significantly reduced the LVEF on day 9. At all three time points, the LVFSs after treatment with AZM (both AZM and HCQ + AZM) showed similar changes with LVEFs. Compared with CON, HCQ treatment did not change the LVEF and LVFS on days 3, 6, and 9. These results demonstrate that AZM and HCQ + AZM can severely alter cardiac function, which are consistent with our ECG findings showing that AZM and HCQ + AZM prolong the QTc interval.

**Figure 5.**
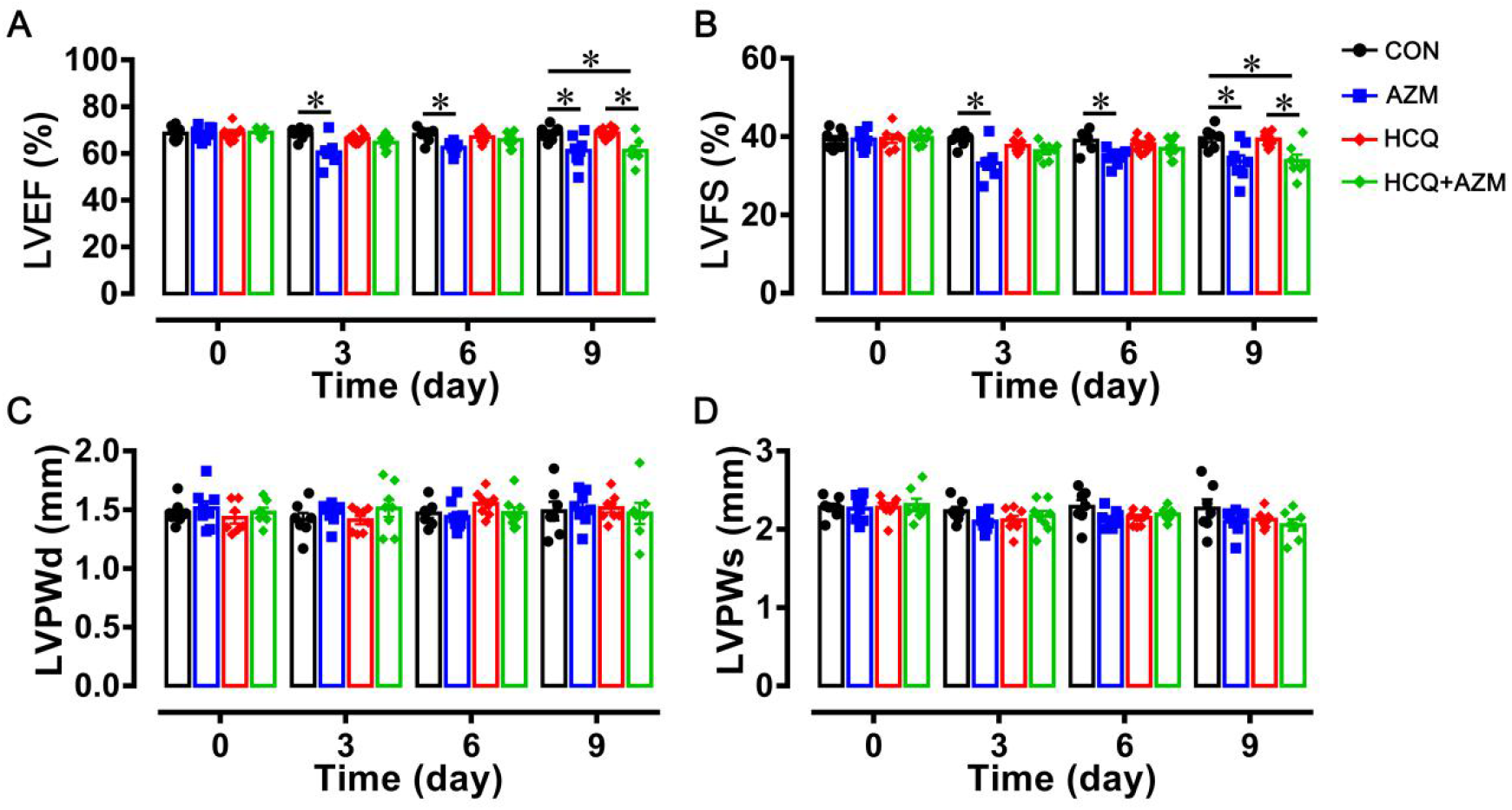
*In vivo* echocardiographic data obtained before drug administration and after 3, 6, and 9 days of hydroxychloroquine (HCQ) treatment with or without azithromycin (AZM). Changes in the ejection fraction (A), fraction shortening (B), LVPWd (C), and LVPWs (D). Data are presented as mean ± standard error of mean (n = 7-8, both groups), *p < 0.05. Statistical significance was determined using two-way analysis of variance coupled with Tukey’s multiple comparison test. CON, control group; AZM, Azithromycin group; HCQ, Hydroxychloroquine sulfate group; HCQ + AZM, Hydroxychloroquine sulfate + Azithromycin group; LVEF, left ventricular ejection Fraction; LVFS, left ventricular fractional shortening; LVPWd, left ventricular posterior wall thickness in diastole; LVPWs, left ventricular posterior wall thickness in systole.

**Table 4.** Echocardiographic parameters of guinea pigs before and after drug administration. Data are presented as mean ± standard error of mean (n = 7–8, both groups). Statistical significance was determined using two-way analysis of variance coupled with Tukey’s multiple comparison test. ^a^P< 0.05 vs. CON group; ^b^P<0.05 vs. AZM group; ^c^P < 0.05 vs. HCQ group. CON, control group; AZM, Azithromycin group; HCQ, Hydroxychloroquine sulfate group; HCQ + AZM, Hydroxychloroquine sulfate + Azithromycin group; D, day; LVPWd, left ventricular posterior wall thickness in diastole; LVPWs, left ventricular posterior wall thickness in systole; LVIDd, left ventricular internal dimension in diastole; LVIDs, left ventricular internal dimension in systole; EDV, left ventricular end diastolic volume; ESV, left ventricular end systolic volume; LVEF, left ventricular ejection fraction; LVFS, left ventricular fractional shortening.

| Time | Group | LVPWd<br>(mm) | LVPWs<br>(mm) | LVIDd<br>(mm) | LVIDs<br>(mm) | EDV<br>(μL) | ESV<br>(μL) | LVEF% | LVFS% |
| --- | --- | --- | --- | --- | --- | --- | --- | --- | --- |
| D0 | CON | 1.48±0.03 | 2.28±0.04 | 6.64±0.22 | 4.05±0.16 | 228.66±17.53 | 73.23±7.33 | 68.44±1.10 | 39.25±0.88 |
| | AZM | 1.51±0.06 | 2.27±0.06 | 6.33±0.13 | 3.87±0.14 | 212.68±3.42 | 70.20±3.00 | 68.54±1.05 | 39.29±0.83 |
| | HCQ | 1.43±0.05 | 2.28±0.06 | 6.56±0.16 | 3.99±0.11 | 221.64±12.04 | 73.97±3.23 | 68.65±1.30 | 39.44±1.09 |
| | HCQ + AZM | 1.48±0.04 | 2.31±0.08 | 6.57±0.20 | 3.97±0.14 | 223.70±15.18 | 69.92±6.03 | 68.92±0.73 | 39.64±0.64 |
| D3 | CON | 1.42±0.06 | 2.23±0.05 | 7.11±0.16 | 4.31±0.16 | 265.30±13.25 | 84.71±7.74 | 68.45±0.94 | 39.45±0.94 |
| | AZM | 1.47±0.03 | 2.10±0.04 | 6.45±0.19 | 4.22±0.15 | 213.89±14.53 | 80.49±6.88 | 60.47±1.90<sup>a</sup> | 33.14±1.41<sup>a</sup> |
| | HCQ | 1.41±0.04 | 2.12±0.05 | 6.62±0.17 | 4.21±0.11 | 226.31±12.91 | 79.69±5.06 | 66.38±0.80 | 37.66±0.65 |
| | HCQ + AZM | 1.51±0.07 | 2.17±0.07 | 6.70±0.16 | 4.38±0.12 | 232.87±12.60 | 87.58±5.77 | 64.57±1.08 | 36.26±0.79 |
| D6 | CON | 1.47±0.04 | 2.29±0.08 | 6.94±0.22 | 4.30±0.13 | 252.54±19.03 | 83.65±6.12 | 67.95±1.17 | 38.99±0.96 |
| | AZM | 1.44±0.04 | 2.12±0.04 | 6.94±0.14 | 4.54±0.07 | 240.07±8.68 | 91.63±2.28 | 62.50±0.89<sup>a</sup> | 34.77±0.70<sup>a</sup> |
| | HCQ | 1.55±0.04 | 2.15±0.03 | 6.54±0.12 | 4.03±0.10 | 219.65±9.30 | 71.60±4.10 | 66.93±0.96 | 32.02±0.73 |
| | HCQ + AZM | 1.47±0.05 | 2.19±0.03 | 6.22±0.17<sup>ab</sup> | 3.98±0.13<sup>b</sup> | 196.72±12.25<sup>a</sup> | 69.63±5.07 | 65.71±1.15 | 36.92±0.91 |
| D9 | CON | 1.49±0.08 | 2.27±0.12 | 7.02±0.27 | 4.39±0.18 | 259.69±23.18 | 88.30±8.83 | 68.61±1.20 | 39.59±1.01 |
| | AZM | 1.51±0.05 | 2.10±0.06 | 6.32±0.21<sup>a</sup> | 4.29±0.20 | 204.69±14.95<sup>a</sup> | 84.62±9.02 | 61.09±2.24<sup>a</sup> | 33.60±1.58<sup>a</sup> |
| | HCQ | 1.52±0.04 | 2.13±0.04 | 6.34±0.14 | 3.82±0.10 | 203.72±11.14<sup>a</sup> | 64.75±4.17 | 68.69±0.85 | 39.29±0.70 |
| | HCQ + AZM | 1.47±0.09 | 2.06±0.07 | 6.67±0.27 | 4.45±0.19<sup>c</sup> | 232.22±21.80 | 91.64±9.40<sup>c</sup> | 61.15±2.09<sup>ac</sup> | 33.76±1.58<sup>ac</sup> |

## Discussion

In this study, we have demonstrated that the combination of HCQ and AZM treatment increase the risk of cardiac dysfunction compared with HCQ treatment. These findings in the healthy animal model is similar with that in the COVID-19 patients treated with this combination. It has been reported that the COVID-19 patients treated with HCQ + AZM treatment for 15 days are associated with higher frequencies of QT interval prolongation(4, 5). Moreover, our results showed that HCQ did not alter the QT interval after short-course treatment in the healthy animals, which is consistent with previous finding indicating that the risk of cardiomyopathy following short-term use of HCQ was low(19). However, some of the COVID-19 patients treated with HCQ have showed prolonged QT interval(4, 5). This might indicate that the cardiovascular risk of HCQ is increased in the COVID-19 patients suffering cardiac injury. Autopsies(10) and experiments(20) have shown that viruses can enter cardiomyocytes, causing myofibril damage. Heart inflammation or myocarditis were also seen in the COVID-19 patients with mild symptoms(20). Thus, we suggest that disease models resembling SARS-CoV-2 induced cardiac damage are needed when assessing the cardiotoxic effects of HCQ.

Drug-induced cardiotoxicity is a major adverse event associated with numerous clinically important drugs. Cardiotoxicity has previously led to the post-marketing withdrawal of numerous pharmacologically active drugs. As a consequence, the assessment of cardiotoxicity potential is a crucial parameter in drug development. Here, in addition to ECG, we used echocardiography to evaluate cardiac function following AZM, HCQ, and HCQ + AZM administration. Currently, echocardiography clinically is an essential tool for testing drug-induced left ventricular systolic dysfunction with a fall in left ventricular ejection fraction(18), but this technology has not been widely used to assess potential drug cardiotoxicities preclinically. On the other hand, it has been reported that patients with long QT syndrome are associated with delayed systolic contraction velocity and prolonged systolic duration(21), and in the patients with LVFS less than 35%, the QT interval is correlated with left ventricular systolic dysfunction(9). Here, we have demonstrated that cardiac dysfunction is consistent with the prolongation of QT intervals in the guinea pigs following either AZM or the combination of AZM and HCQ treatment. Thus, our and others’ results suggest that echocardiography technology are useful for assessing the cardiotoxicity of AZM and HCQ. Another advantage of echocardiography is that it provides morphological measurements of the heart. Thus, without sacrificing the animals, we can detect any potential structural changes induced by newly developed drugs.

## Limitations

This study has just explored the effects of AZM and the combination of HCQ + AZM on cardiac conduction and function at a single dose. In the coming study, a higher and a lower dose have been selected to unveil the other potential hidden cardiotoxicity. The protein expression alterations of cellular membrane receptors and their downstream signals are to be investigated to explain underlying molecular mechanisms.

## Author Contributions

F. H. Yang, Y. Zhang, X. Li designed and initiated the project. X. Li, S. Zheng, W. Tan, H. Sun, X. Zhang, X. Li, H. Chen, X. Ren, T. He, were responsible for the laboratory experiments, data analysis, and/ or animal care. All authors read and approved the final manuscript.

